# A shared neural substrate for action verbs and observed actions in human posterior parietal cortex

**DOI:** 10.1101/2020.04.20.039529

**Authors:** T. Aflalo, C. Zhang, E.R. Rosario, N. Pouratian, G.A. Orban, R.A. Andersen

## Abstract

High-level sensory and motor cortical areas are activated when processing the meaning of language, but it is unknown whether, and how, words share a neural substrate with corresponding sensorimotor representations. We recorded from single neurons in human posterior parietal cortex (PPC) while participants viewed action verbs and corresponding action videos from multiple views. We find that PPC neurons exhibit a common neural substrate for action verbs and observed actions. Further, videos were encoded with mixtures of invariant and idiosyncratic responses across views. Action verbs elicited selective responses from a fraction of these invariant and idiosyncratic neurons, without preference, thus associating with a statistical sampling of the diverse sensory representations related to the corresponding action concept. Controls indicated the results are not the product of visual imagery nor arbitrary learned associations. Our results suggest that language may activate the consolidated visual experience of the reader.

## Introduction

How do words get their meaning? Although the exact architecture of the semantic system is still under debate, most evidence suggests that meaning emerges from interactions between supramodal association regions that code abstracted symbolic representations and the distributed network of regions that process higher-level aspects of sensory stimuli, motor intentions, valence, and internal body-state (*1–5*). Engagement of the distributed network is taken as evidence that the brain’s representation of the physical manifestation of words is an important component of their meaning. For example, visual coding for the form of a banana, the motor act of biting into or peeling a banana, and its taste and texture would be components of meaning in addition to more symbolic, lexical aspects of meaning such as the “dictionary definition”. Although this view is generally accepted, no single unit recording evidence has demonstrated a shared neural substrate between processing the meaning of a word and its visuomotor attributes within the distributed network. To date, supporting evidence comes from lesion and fMRI studies establishing a rough spatial correspondence between brain areas involved in high-level sensorimotor processing and areas recruited when reading text or performing other behaviors that require access to meaning (*1, 6*). A lack of direct neural evidence is concerning given that neuroimaging (*7*) and lesion results (*8–10*) have been mixed, and cannot establish a shared neural substrate at the level of single neurons (*11*). Thus how words get their meaning translates into two immediate questions with regard to single neuron selectivity: one, are words and their sensorimotor representations coded within the same region of cortex; and, two, is there a link between words and their sensorimotor representations? In this paper, linking will refer to the existence of a shared neural substrate with individual neurons exhibiting matching selectivity for both a word and the corresponding visual reality.

To complicate matters, the number of sensorimotor representations that can be described by the same basic concrete word are generally very large (e.g. the visual form of a “banana” depends on ripeness, viewing angle, lighting, and whether it is peeled or sliced) and invariance is very rarely complete in high-level sensorimotor regions (*12–14*). This raises a third question: If the same object is coded in different ways depending on details of presentation, how might a word link to these varied visual representations? Stated more generally: What is the neural architecture that links neuronal responses to silently reading a word and seeing varied visual presentations of what the word signifies? The answer is critical in understanding how sensorimotor representations influence our understanding of words. Do we connect the symbolic representation of a word to an abstracted invariant and therefore universal visual representation? To a particular canonical example? Or to the many diverse representations that comprise our varied experiences? The question applies to all concrete words that describe physical reality, including action verbs. In this study we look at how neural coding for action verbs relates to varied visual representations of corresponding observed actions.

Finally, what cognitive phenomena can account for the presence of a link between a word and its visual representation within any experimental paradigm? The link may mediate semantic memory, reflecting associations between the word and its visual representations built over a lifetime of experience. In this view, reading words activates sensorimotor representations automatically and these representations are an intrinsic *component* of the meaning of the word. Second, reading a word has been hypothesized to evoke mental imagery. Responses in sensorimotor cortex may reflect such imagery and the link could be between visual representations and mental imagery of the same stimuli. Or the link may be the consequence of short term learning such as occurs during categorization (*15*). Given these multiple possibilities we address a fourth question: if a link exists, what cognitive process does the link mediate?

To address the above four questions, we recorded populations of neurons from electrode arrays implanted in two tetraplegic individuals (NS and EGS) participating in a brain-machine interface (BMI) clinical trial while the participants viewed videos of manipulative actions or silently read corresponding action verbs. The implants were placed at the anterior portion of the intraparietal sulcus (see Fig S1 for implant locations), a region that is part of the “action observation network” (AON) composed of the lateral occipital temporal cortex (LOTC, (*16*)) as well as frontal and parietal motor planning circuits (*17, 18*). These regions are involved in higher-order processing of observed actions (*19–22*) and neuroimaging and lesion evidence implicate a role in verb processing (*23–29*). The ability to perform invasive neural recordings provides us with the first opportunity to probe whether and how language links with corresponding visual representations at the level of single neurons in high-order sensory-motor cortex. Towards this objective we establish four primary results relating to the four questions outlined above: First, PPC neurons show selectivity for action words and visually observed actions; Second, a portion of PPC neurons link action verbs and corresponding visual representations; Third, text selective units in PPC link with all the diverse visual representations found in the neural population; and, fourth, the link is not based on imagery or short-term learning and thus appears to be semantic in nature. One possible interpretation is that when reading text, we replay our visual history as part of the process of understanding and thus ground our conceptual understanding in our unique experiences.

## Results

Participants viewed videos of five manipulative actions presented in 3 visual formats (two lateral views differing in body posture and one frontal view) and a fourth format, text, requiring the subject to silently read associated action verbs (See Fig 1A for example stimuli). Five actions were used: drag, drop, grasp, push and rotate, for which preliminary experiments (Fig S2) had demonstrated neuronal selectivity. A total of 15 unique videos (five actions by three visual formats) and 5 written action verbs were presented. Presenting the observed actions in three formats allowed us to tease apart different models of how action verbs associate with overlapping (common to visual formats / exhibiting invariance across all formats) and distinct (idiosyncratic to given formats / not invariant or only invariant across subset of formats) features of the neural code for observed actions. This design allowed us to answer the first three questions posed in the introduction. We recorded 1586 units during 18 recording sessions in two subjects. For the first 7 sessions in subject NS, and all sessions for subject EGS, the participants passively watched the action videos and silently read the action verbs. Results from silent reading and active imagery were quantitatively similar in NS and thus data was pooled across sessions when addressing the first three questions of this paper. To answer the fourth question, for the final 6 sessions in subject NS, the participant used the action verb as a prompt to “replay” the associated action video using visual imagery from either frontal (F) or lateral (L0) perspectives, thus allowing us to quantify how imagery affects verb processing. Additionally, for question 4, we present a control study in which abstract symbols are paired with visual imagery of motor actions to better understand the effects of short-term associations.

**Figure 1:**
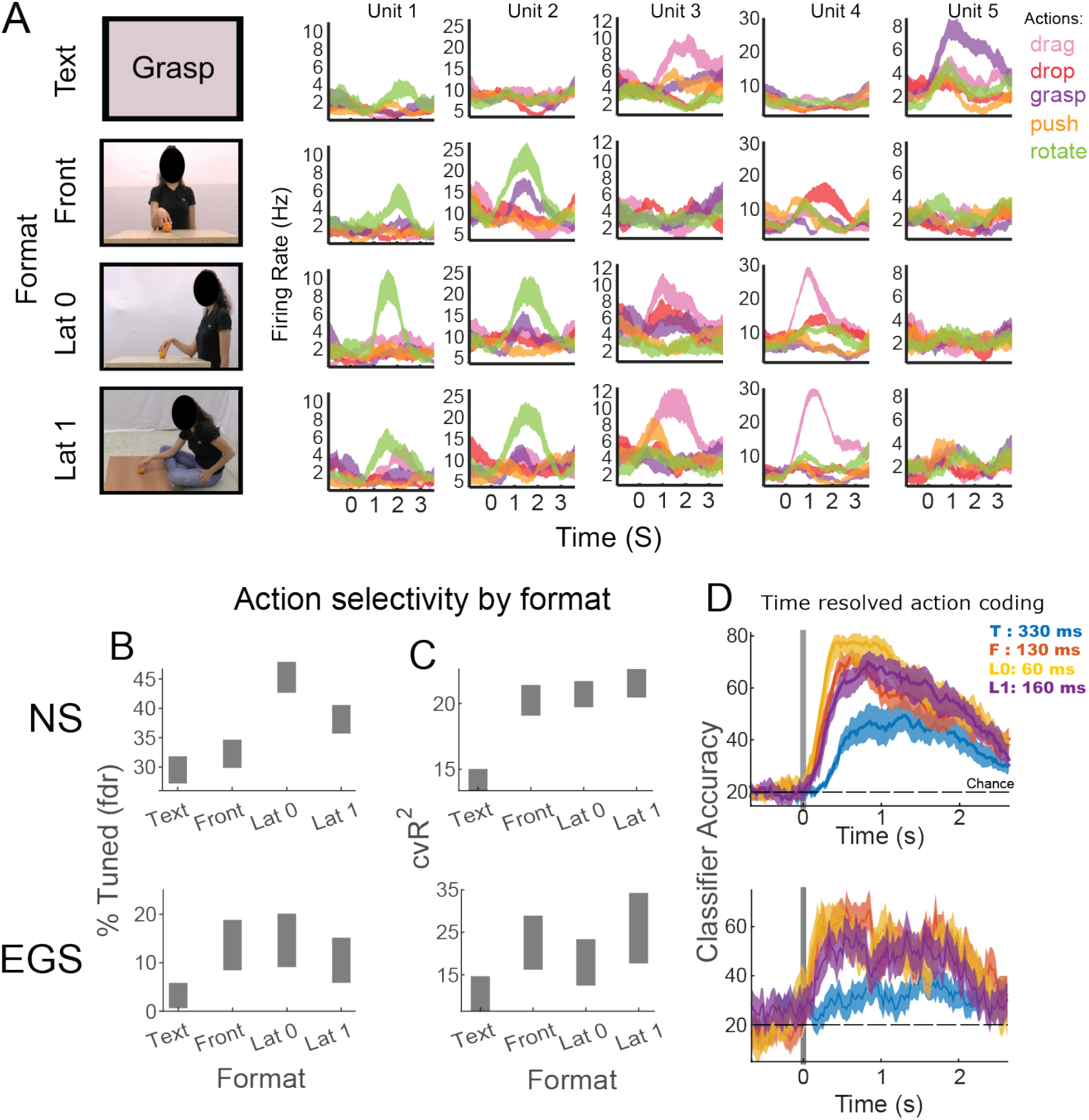
Human parietal neurons are selective for observed actions and action verbs. **(A)**. Example neurons illustrating diverse selectivity patterns across formats. *Left panel*: Sample still frames depicting “grasp” in each format. Face is masked to obscure identity per publisher’s request. *Right panel*: Representative units illustrating diverse neural responses to the five tested actions (color coded) across the four tested formats. Each panel shows the firing rate (mean ± sem) through time for each action for a single format. Each column illustrates the responses of the same unit to the four formats. See figure S1 for recording locations. **(B)** Percent of units with significant action selectivity split by format (mean ± 95% CI, one-way ANOVA, p<0.05 fdr corrected.) Zero units were selective in each format during the 1 second window prior to stimulus onset (one-way ANOVA, p<0.05 fdr corrected) **(C)** Cross-validated R^2^ of units with significant selectivity (units significant in B) split by format (mean ± 95%CI). **(D)** Sliding-window within-format classification accuracy for manipulative actions. Sliding window= Overlapping 300ms windows with 10ms increments. Inset displays color code for format and associated latency estimate for onset of significant decoding.

### Are human posterior parietal cortex (PPC) neurons selective for observed actions and action verbs?

Figure 1A shows the response of five representative neurons illustrating the variety of selectivity for both observed actions and action verbs at the level of individual neurons. Within a format, we defined units as selective if there was a significant differential response, measured as average firing rate over video duration, to the different action identities. Significant selectivity across the recorded neural population was found for action verbs, however, the percent of selective units as well as the consistency of the response, as measured by the cross-validated coefficient of determination (cvR^2^), was smaller for text than for observed actions (Fig 1B, C). All five actions evoked significant neural responses from baseline across the four formats (Fig S3). The majority of visually selective units were driven by the video presentations, as in non-human primate anterior intraparietal area (AIP(*22*)). A minority, however, were suppressed by the video and text presentations (Fig S3). The mean response strength decreased smoothly from the action evoking the maximal response to the weakest response. Individual units could show steep or more graded selectivity, and this pattern was essentially identical across formats (Fig S4). Greater selectivity for action videos relative to text was reflected in a time-resolved decode analyses (Fig 1D). Defining the latency of action selectivity as the onset of significant classification accuracy revealed shorter latencies for the visual formats (windows starting at 60 −160ms depending on format) than the written word (330ms), possibly reflecting differences in afferent pathways. Taken together, all formats were encoded within the population, but with greater selectivity and shorter latency for videos relative to text.

### Is there a link between neural representations of action verbs and observed actions in human PPC?

Having established that PPC neurons are selective for both action verbs and observed actions, we now ask whether there exists a shared neural substrate, with neurons exhibiting matching selectivity for both a word and the corresponding visual representation. We addressed this by using two population analyses: across-format classification and across-format correlation. Leave-one out cross-validation was used to train a classifier to predict action identity within-format. On each fold, the decoder was also used to predict action identity from the three additional formats. This across-format generalization analysis measures how well the neural population structure that defines action identity in one format generalizes to other formats (Fig 2A, B). Across-format accuracy was above chance for all pairs of formats for NS and above chance when pooling across visual formats to achieve adequate power for EGS (Rank-sum test, p<0.05). In contrast, across-format accuracy was not significant when shuffling action identity between formats (Shuffled Alignment, red in Fig 2A, B). This result demonstrates that the neuronal representation was not random; the population is more likely to link representations across formats for the same action identities. However, the results also demonstrate that the generalization is not perfect: the across-format accuracy is lower than the within-format accuracy suggesting that the neural code for action identity also depends on details of presentation. The strength of generalization was format dependent being near perfect across body-postures (same lateral view), still high, but reduced across shifts in viewing perspective (across the lateral and frontal views), and lowest when comparing observed actions with the written verb.

**Figure 2:**
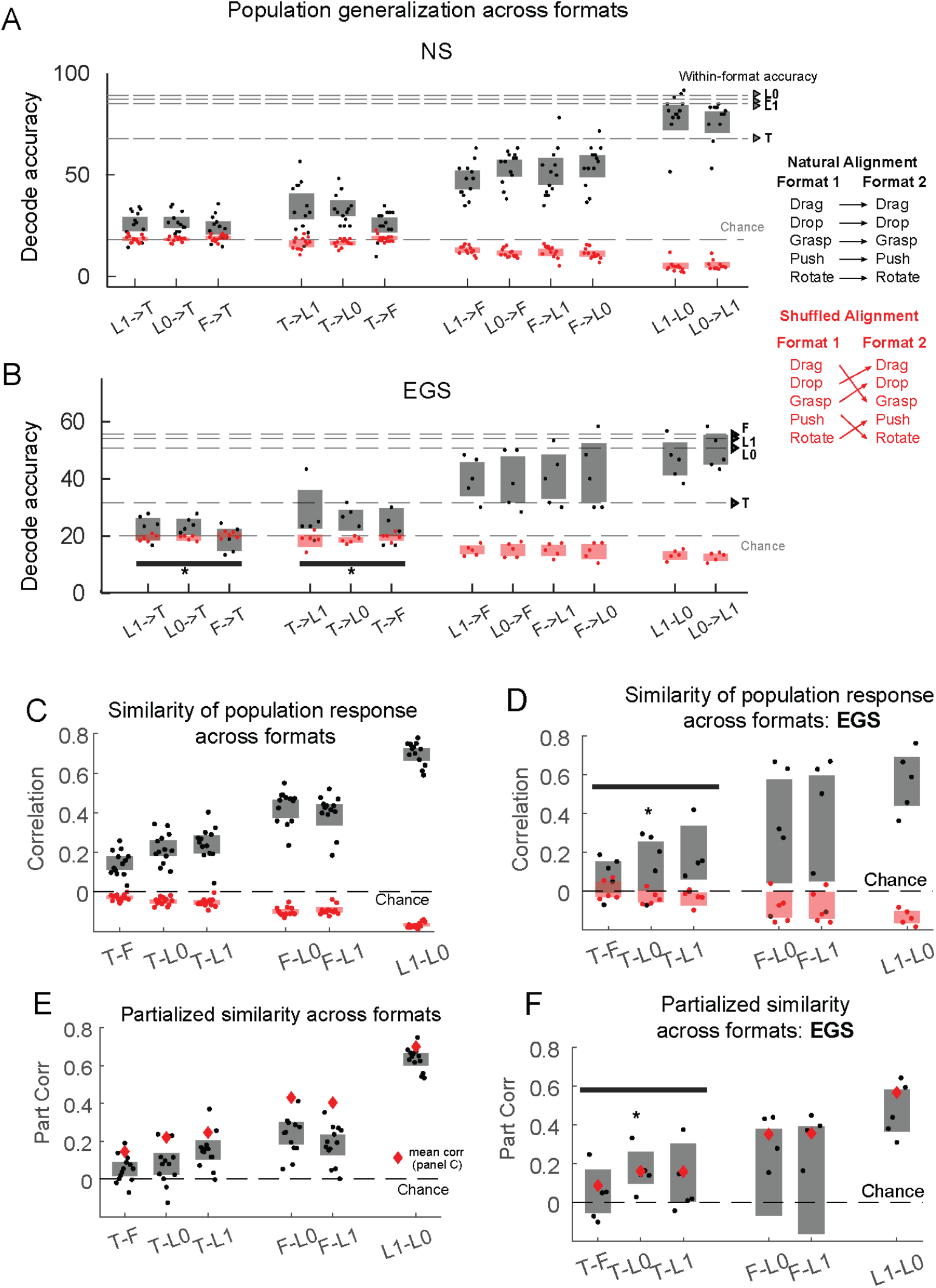
Action verbs link with observed actions. **(A)** Across-format classification of manipulative actions. X-axis labels indicate the formats used for classifier training and testing (e.g. train->test.) Dots = single session result. Rectangle = 95% bootstrapped CI. Grey (red): values for matched (mismatched) labels across formats (see inset for definitions). Dashed horizontal lines show within-format cross-validated accuracy (mean across single session results.) All comparisons with chance performance (dashed line) reached significance (Wilcoxson rank sum test, p<0.05). **(B)** Similar to A but for EGS. Cross-format classification significant between all visual formats, and between visual and text formats when pooling visual formats (see bar with asterisk.) **(C)** Correlation of neural population responses across pairs of formats. Conventions as in A. **(D)** Same as D for participant EGS. **(E)** Pairwise population correlation while controlling for additional formats using partial correlation. Resulting correlations are above chance (part corr =0) but below standard correlation values (mean= red diamonds.) **(F)** Same as F for participant EGS

Significant generalization of action representations across formats was robust to the analysis technique. We correlated neural population responses across formats (Fig 2C, D). Population responses were constructed by concatenating the mean response of all units to each action within-format (Fig S5). A significant positive correlation was found for all format pairs while no significant positive correlation was found when shuffling action identity between formats. One caveat to interpretation is that the correlation between any pair of formats may be the consequence of the two formats being correlated with a third format. A significant link between pairs of formats was preserved but somewhat reduced when controlling for the other formats using a partial correlation analysis (Fig 2E&F). This last result indicates that text links with each of the visual formats directly as the significant link is preserved when the possible mediating factors of the other formats are removed.

The preceding population analyses established that text and visual representations are linked pairwise at the level of the population, but the link does not perfectly generalize across formats. What is the breakdown of the single units that compose the population results? To answer this question, we compared the precise selectivity pattern (SP, defined as the firing rate values for each of the five actions) across pairs of formats using a model selection analysis for each neuron. A linear tuning model can describe the 4 possible ways the SP can compare across two formats (Fig 3A). **(1)** Both formats are selective in a similar manner (Fig 3A, Matched selectivity); the linear parameters (***α**ϵR*^5^) for each of the five actions are constrained to be identical for the two formats. **(2)** Both formats are selective, but with mismatched patterns (Fig 3A, Mismatched selectivity) the linear parameters (***α, γ**ϵR*^5^) are different between the two formats. **(3, 4)** Finally, only one of the two formats may be selective (Fig 3A, Single format-1or format-2 selective); a constant scalar offset term is used for the non-selective format (not shown). We identified the model that best described the neuronal behavior using both the Bayesian information criteria (BIC) and the cross-validated coefficient of determination (cvR^2^). We found that the two measures provide complementary perspectives when comparing across formats (Fig S6.) In summarizing the results, we used the average percentages provided by both measures. In line with our population results, we found that the percent of cells with a similar selectivity patterns across formats (Fig. 3B&C, red) was format-dependent, being greatest across body-postures (same lateral view), slightly reduced across shifts in viewing perspective (across the lateral and frontal views), and lowest when comparing observed actions with the written verb. These results not only indicate that text links with the visual formats and the visual formats link with each other, but also that a percentage of the population codes the same action identities in different formats with differing patterns of selectivity.

**Figure 3:**
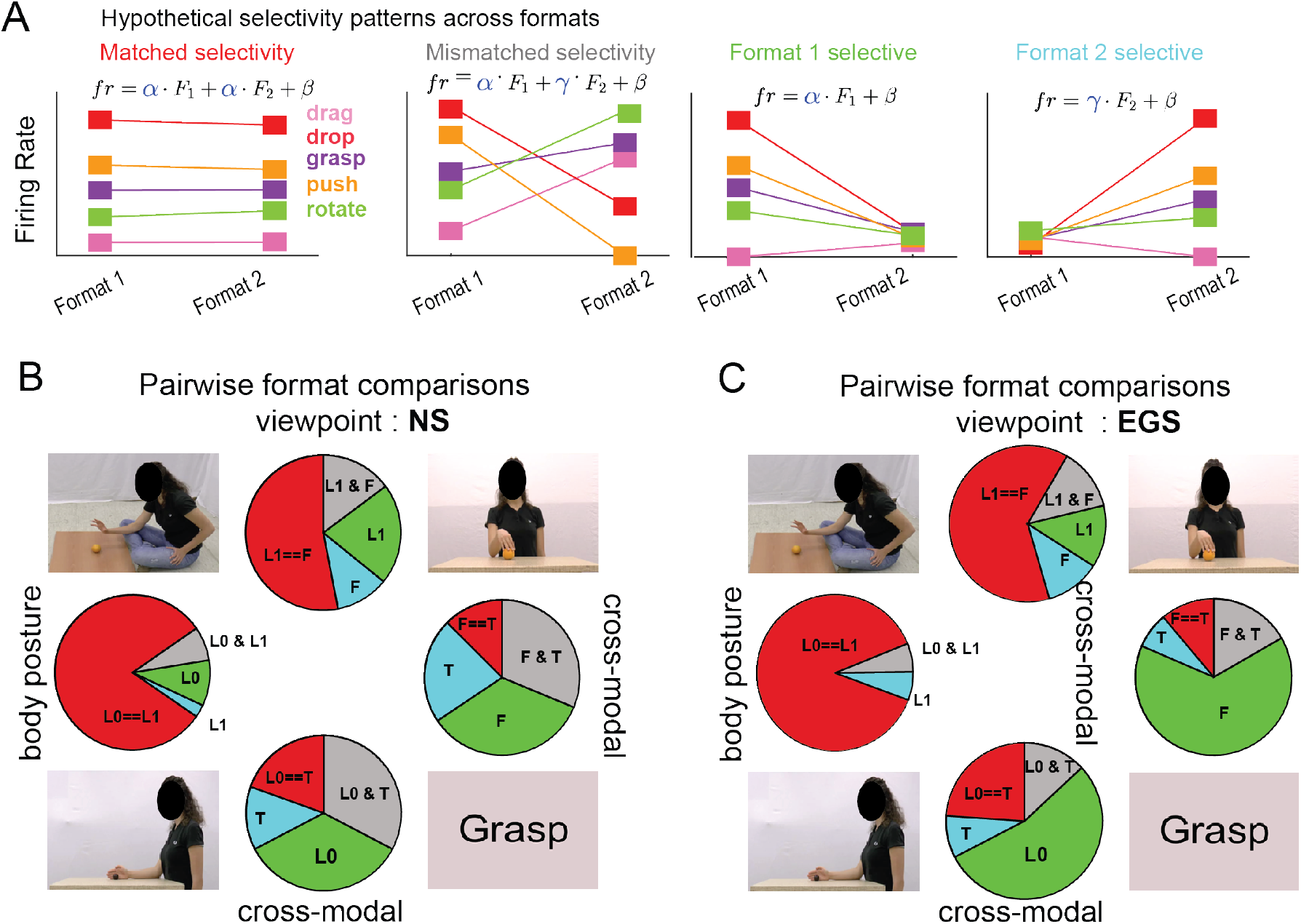
Single neuron selectivity patterns link action verbs and observed actions. **(A)** Schematic illustrating the four possible ways the selectivity pattern (SP) can compare across two formats (see Fig S6 for expanded description). **(B)** Summary of SPs across pairs of formats for participant NS (see Fig S6). Red = matched SP, grey = mismatched SP, cyan, light green= selectivity for a single format only (see title colors in panel A). Face is masked to obscure identity per publisher’s request. **(C)** Same as panel B for participant EGS.

### What is the architecture that links observed actions and action verbs?

The preceding section demonstrated that there is a neural link between action verbs and visually observed actions. Here we seek to understand the architecture of this link: to characterize how text-selective units link with the varied visual presentations of the same action. As a prerequisite, we first characterized how the different visual presentations were encoded with respect to each other, ignoring the text format. Just as neural selectivity patterns can compare across two formats in four different ways (Fig 3A), they can compare across 3 formats in 14 possible ways (see Fig. 4A, X-axis labels and examples). As above, a model selection analysis was used to categorize each unit based on the model that best described the selectivity patterns across the visual formats (Fig 4A). The population was heterogeneous, characterized by units with matched SPs and mismatched SPs in varied combinations across the different visual formats. This diversity can be seen in the individual unit examples of figure 1A; units 1 and 2 shows matching patterns of selectivity across all the visual formats (Fig 4A, L0=L1=F), unit 3 shows matching selectivity across two of the visual formats and no selectivity in the third (Fig 4A, L0=L1), and, unit 4 shows matching selectivity between two formats and mismatching selectivity in the third (L0=L1&F). Thus we find that presentation details impact neural coding for action identity and that individual units link action identity across formats in an assortment of ways when considering all three of the visual formats at once. This result is consistent with the significant but incomplete generalization of action identities across the visual formats shown in figures 2 and 3.

**Figure 4:**
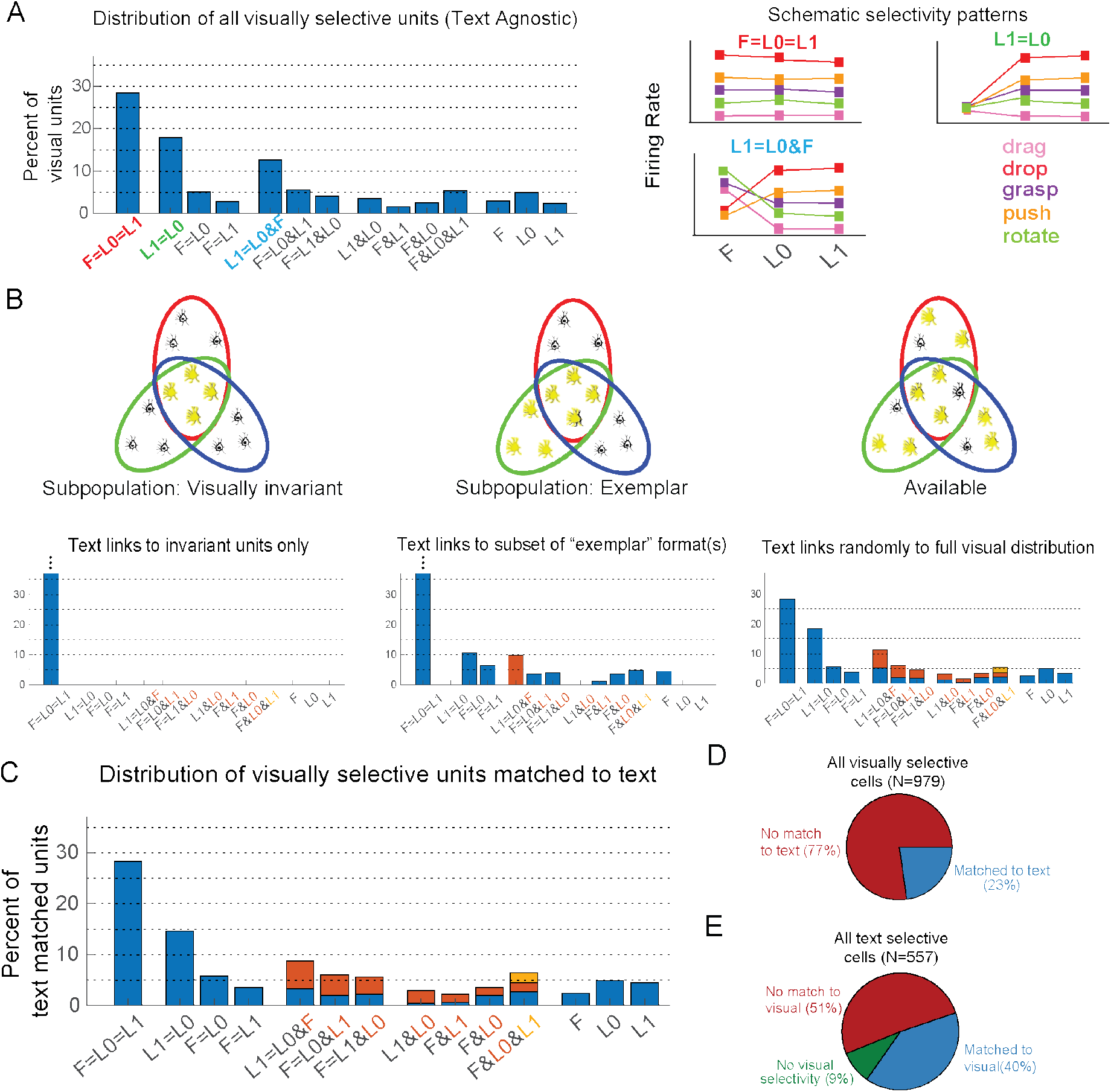
Text links with all available visually selective cells. **(A)** Histogram characterizing how the population of neurons link action representations across the three visual formats (F,L0,L1). “=” matched selectivity pattern (SP), “&” mismatched SP. Exclusion of a format indicates no selectivity. Three schematic selectivity patterns (right panel, color coded) across the visual formats are shown to illustrate how the SPs compare across formats. **(B)** Schematic models illustrating different architectures of how text relates to 3 visual representations of the corresponding action. Each oval contains the population of neurons that are selective for a particular visual format. Overlap between ovals indicates matching selectivity across formats. The possible patterns of overlap between ovals may be more complicated (e.g. more overlap between 2 of the three ovals) but is simplified here for schematic purposes. Yellow neurons are selective for text with matching selectivity while grey neurons are not. Underneath each schematic, is a prediction for how the distribution in panel A will change when the model selection analysis filters the full distribution of A for units with matching text selectivity. **(C)** Similar to A, however the histogram is limited to the subset of visually selective units with a matched SP to text (blue subpopulation in panel D). In cases where the units have mismatched visual SPs (e.g. L0 & F), text can have a matched SP with one of several of the visual formats. Colored segments of histogram indicate which format has matched SP with text (see x-axis labels for color code). **(D)** Percentage of visually selective units with a matched SP to text. **(E)** Percentage of text selective units with a matched SP to at least one visual format, mismatched SP to visual formats, or without visual format selectivity.

Having established that the same action is coded in different ways depending on details of visual presentation, we can now look at how action verbs link to these varied visual representations. We can frame our question in the following way: do action verbs link with the entire population of cells demonstrating visual selectivity or specific subpopulations of cells? Figure 4B illustrates these possibilities. Two primary theoretical possibilities in the literature describe how text can link with subpopulations of visually selective neurons. *Overlapping* describes the architecture in which verbs link specifically with the subpopulation of neurons that are invariant across the visual formats(*30*). *Exemplar* describes the architecture in which verbs link with a specific prototypical exemplar, or “best example”, of the word (*30*). The exemplar may be of a single visual presentation or some subset of presentations.

Finally, we term the situation in which text links with all visually selective cells as *Available*. In this architecture, the link between text and the visual representations mirror the statistics for how the visual representations are encoded within the neural population independent of text. Underneath each schematic, we provide a prediction for how the distribution of figure 4A should change when the smodel selection analysis accounts for how text links with the visual formats.

We extended the model selection analysis to categorize each unit based on the model that best described the selectivity patterns across all four formats (text + all visual formats.) We compared the distribution of the visually selective units with a matched SP to text (Fig 4C) to the full distribution of the visually selective units (Fig 4A). The distribution was essentially unchanged; the subset of visually selective units that link with text reflects a random sampling of the visually selective units: a bootstrapped correlation analysis comparing the empirical distribution of figure 4C with the predictions of 4B show that the population best matches the Available model (correlation with invariant =0.32, exemplar=0.48, available=0.97). This provides the answer to the question of architecture: the distribution of text-linked units (Fig 4C) mirrors the statistics of how visual formats are encoded independent of text, or, in other words, text forms links with all available visual representations. Units with a matching SP between text and at least one visual format (the distribution of 4C,) represents 23% of all visually selective units (Fig. 4D) and 40% of all text selective units (Fig. 4E).

### What cognitive process does the link between action verbs and observed actions mediate?

Does the link between text and the visual formats reflect a semantic association, visual imagery, or short-term, learned associations that formed through the course of the experiment? Thus far, our analyses are based on averaging the neural response across the video duration. This large temporal window may encompass multiple cognitive processes. If neural processing for action verbs specifically reflects bottom-up semantic processing, we would expect to find a shared neural response between formats very soon after stimulus presentation. To address this issue we performed a dynamic, sliding-window cross-validated correlation analysis to look at how the relationship within- and across formats evolves in time (Fig. 5A, B.) To understand how quickly the correlation between text and the visual formats emerges, the diagonal elements of the dynamic correlation matrices were extracted and plotted together for direct comparison in the inset panels of figure 5A, B. These results show that the cross-modal link between text and the visual formats is fast: the onset of the cross-format correlation between text and the visual formats is the same as the within-format text correlation. In other words, as soon as a neural response to text emerges, it immediately shares a common activation pattern with the observed actions.

**Figure 5:**
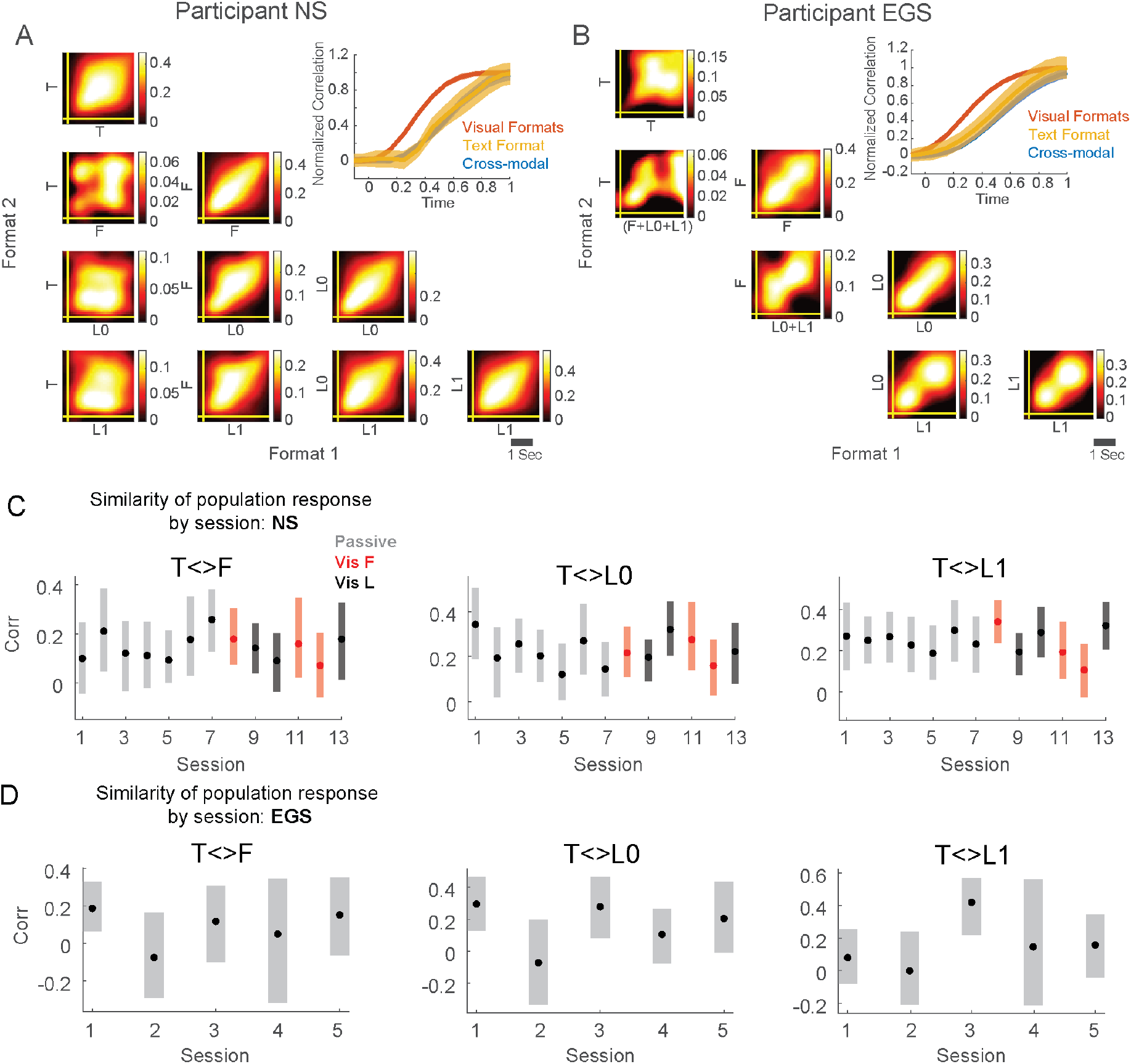
Temporal features support a semantic link between verbs and observed actions. **(A-B)** Cross-modal match between text and visual formats occurs at low latency. **(A)** Dynamic cross-validated cross-correlation matrices demonstrating how the neural population response during stimulus presentation at one slice of time compares to all other slices of time, both within and across formats. Format comparisons as shown in x- and y- axis labels. Correlation magnitude as indicated by color bar. Inset: The diagonal elements of the within- and across format matrices were averaged into three logical groupings (1,within-format visual, 2,within-format text, and 3,across-format text to visual) and normalized to a peak amplitude of 1 for comparison purposes. The temporal profile of the averaged correlations (mean±se across sessions) is plotted to emphasize the similarity of onset timing for the within-format text and across format text to visual population correlations. **(B)** Similar to A, but for participant EGS. To compensate for the smaller number of sessions, we grouped correlation matrices for cross-modal comparisons. **(C,D)** Stable relationship between text and observed actions through experimental sessions. **(C)** Cross-format correlations for subject NS shown for text and the visual formats on a per session basis (mean with 95% bootstrapped CI.) Color code shows whether the subject was passively viewing stimuli or asked to actively imagine from the lateral or frontal perspective (see inset, Vis F= visualize from frontal perspective; Vis L = visualize from the Lateral 0 perspective.) **(D)** Same as panel A except for participant EGS (only silent reading).

Next, we checked whether the strength of population correlation changed over the course of the experiment. If neural processing for action verbs reflects a semantic association, we would expect to find the correlation between text and videos to be present from the first session throughout the course of the experiment. In contrast, if the correlation between text and action videos is a product of learned associations that developed over the course of the study, we would predict that the strength of correlation would increase over the course of repeated exposure to the action videos and text. We found that the early correlation response (cross-validated correlation over 1st second of video presentation) between text and the three visual representations for each session, did not depend on session number (Figure 5C, D), favoring the semantic interpretation.

We performed a number of control analyses and manipulations to address the possibility that associations between text and observed actions reflect visual imagery. In six sessions participant NS was instructed to use visual imagery to “replay” the associated action video in her mind from either the front (F) or side (L0) perspectives when given the action verb prompt. If imagery were a dominant factor in establishing the link between text and observed actions, the explicit manipulation of visualizing from the F or L0 perspective should bias the percent of cells with a matched SP in favor of F or L0. However, both the total number of significant units and the population coding structure were essentially unaffected by the explicit task instruction (Fig. 6 A, B). This result shows that the basic link between actions verbs and observed actions is not dependent on the contents of visual imagery. To probe this result further and ensure the subject followed task instructions, we split the dynamic correlation analysis between the passive and active imagery sessions. Intriguingly, we found (Fig 6C, D) that the early correlation was largely unaffected by the behavioral manipulation while the later (e.g. > 1sec) correlation did show significant differences (paired t-test, p<0.05 on pixel values split between passive and imagery sessions), with active imagery increasing the late period correlation strength relative to passive viewing (Fig. 6E). This result suggests that the subject followed task instructions and that imagery can affect neural responses, but that the early responses of neurons are independent of the contents of imagery.

**Figure 6:**
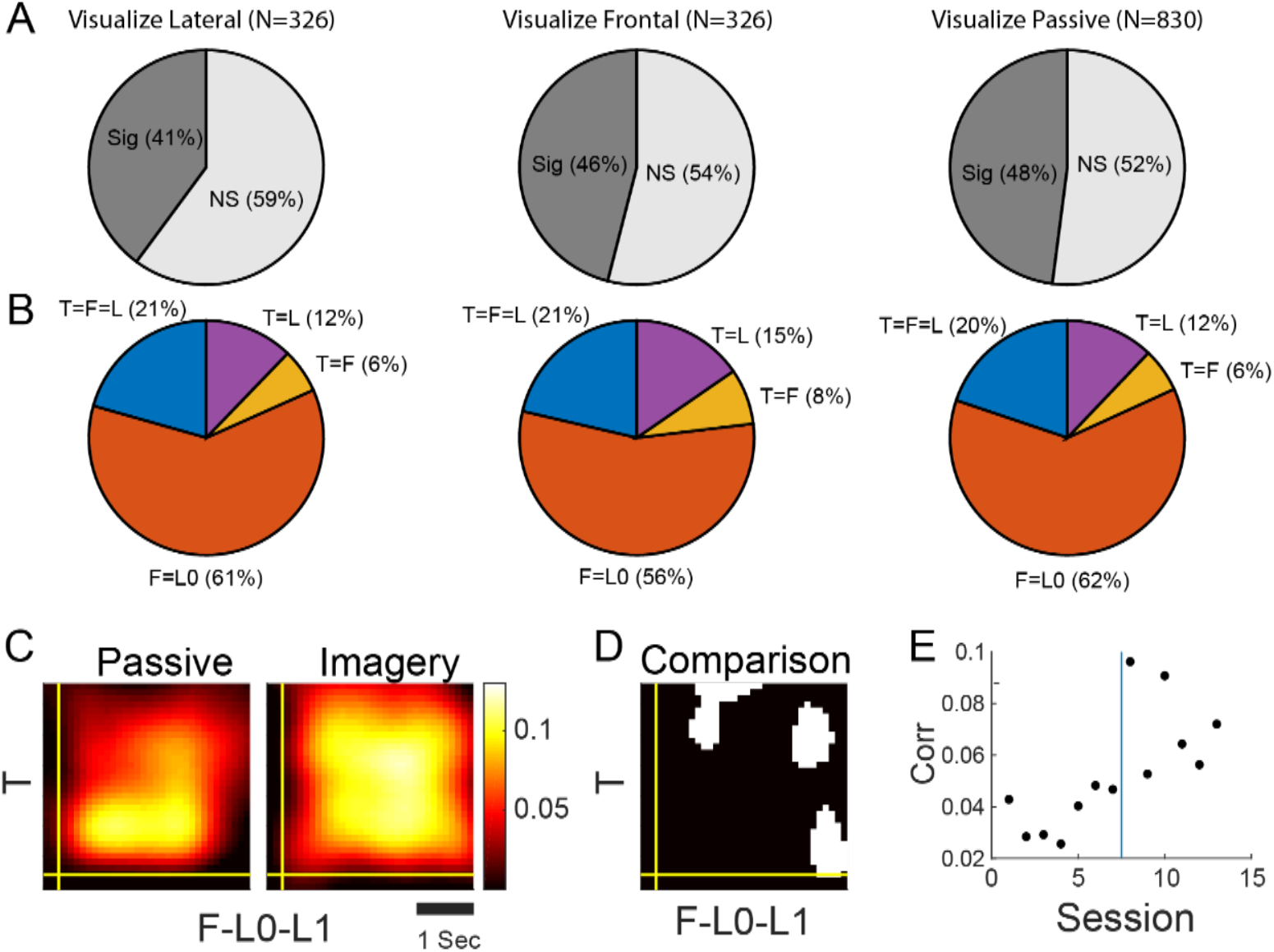
The effect of explicit instruction on cross-format invariance. During the initial seven sessions subject NS silently read action verbs. In the six subsequent runs she explicitly visualized the frontal (F, 3 runs) or lateral standing (L0, 3 runs) perspective in response to the action verb. **(A)** The percent of units with a significant effect of action or action-format interaction for the format by action ANOVA applied to the triplet of formats pertinent to task instruction (T,F,L0). “Sig” = significant at p<0.05 fdr corrected (“NS” otherwise). Results are split by the task instruction. Total number of sorted units shown in title. **(B)** Results for the combined (BIC+cvR2) model selection analyses for the same triplet of actions split by task instruction. The percentage of T=L units was twice as prevalent as T=F units for passive viewing as well as the two instructed conditions. **(C)** Mean dynamic cross-correlation between the visual formats and text split by passive viewing and active imagery in participant NS. **(D)** Pixel coordinates demonstrating a significant difference between passive viewing and active imagery (significant pixels in white, paired t-test, p<0.05.) **(E)** Cross-correlation value between text and the visual formats for the set of significant pixels shown in D as a function of session number. Blue line shows split between passive and active imagery sessions

In a final control, we collected a dataset in which four abstract symbols (snowflakes, Fig S7A) were paired with visual imagery of movements for over 2 months (31 recording sessions, 114 ± 11 units per session, Fig S7B). In this paradigm, subject NS was asked to visualize a movement from the first-person perspective when presented with a symbol. The subject learned this task well, as we could accurately decode the different symbols when the subject was instructed to use visual imagery (Fig S7C). We also asked the subject to passively view the same stimuli at sporadic intervals (Fig S7B, vertical orange lines) and found that the ability to decode the different symbols disappeared (Fig S7C, D). These clear differences in decoding demonstrate that the subject is capable of comprehending and following task instructions as they relate to passive viewing versus active visual imagery. Furthermore, the study shows that not all types of visually distinct stimuli elicit a differential neural response under passive viewing. Finally, it demonstrates that the recorded population does not form automatic neural responses to arbitrary abstract symbols, even when the different symbols have been learned and are of direct behavioral relevance.

## Discussion

Our results answer the four questions raised in the introduction: PPC neurons exhibit selectivity for action verbs as well as observed actions; text links to visual representations of observed action; text links with a fraction of all available visual representations; and, the link is most consistent with being semantic in nature and not due to imagery nor learned associations.

### Answers to the four questions

#### First, selectivity

Both single cell properties and within-format decoding demonstrate neuronal selectivity for action verbs and observed actions in human PPC. The visual selectivity had short latencies (about 100ms), while text selectivity emerged nearly 200ms later. The features of the visual stimuli that determined neural selectivity remains unclear. The term selectivity for action identity should be interpreted as a label assigned to the visual stimuli rather than coding for the basic-level type of action e.g. “grasp”. Manipulations of viewpoint or fixation point (Fig S2) changed neural coding significantly. Manipulative actions can differ in hand and arm postures, contact points with the object, dynamics, amongst others, and these parameters should affect neural coding to represent the behavioral complexity of natural actions. Elaborating the exact degree to which neural coding is influenced by action identity, its many parameters, or even low-level visual features needs further work (see ^34^). Nonetheless, the link between action verbs and observed actions demonstrates that coding of action identity cannot solely be driven by irrelevant visual features. Further, not all visual differences are encoded by the neural population (Fig S2). Finally, high-dimensional coding of both category-relevant and -irrelevant visual features is consistent with neural coding in high-level regions of the ventral visual stream (*12, 31*).

#### Second, action verbs and observed actions share a common neural substrate

We demonstrate the shared substrate at the population level using cross-format decoding and population correlation between formats (Fig. 2), and showed the basis of this population link by modeling of single cell selectivity across pairs of formats (Fig. 3). Prior neuroimaging evidence indicates a degree of anatomical overlap within the AON for processing observed actions and language (*23, 24, 26–28, 32*). However, imaging evidence can be inconsistent (*7*) and gross anatomical overlap seen in neuroimaging does not directly imply that the same neural populations support both tasks (*11*). Our evidence provides definitive evidence for a shared neural substrate by demonstrating that the precise selectivity patterns for action verbs match the selectivity patterns for corresponding observed actions at the neural unit level.

#### Third, architecture

We have established that, at the neural level, action verbs link with visually observed actions, suggesting that sensorimotor representations are an intrinsic *component* of verb meaning. The potential implications of this finding are hard to pin down without understanding the architecture of the link. There are infinite visual stimuli that could be considered a “grasp” or a “banana” or any basic category colloquially used to describe an object or action. Indeed, our results established that neural coding for action verbs depends upon the details of presentation (see Fig 4) and at least some degree of dependence upon presentation details appears to be ubiquitous throughout cortex (14). Given the diversity of neural coding, there are three likely architectures (Fig 5A), each with their own implications for how linking is made between symbolic and visuomotor representations. Text could link exclusively to the subpopulation of cells that are visually invariant across the different visual presentations (e.g. Fig. 4B, “visually invariant”.) In such a case, the aspect of “meaning” conveyed by the sensory-motor representation would be what is universal or common to all presentations. In other-words, sensory-motor meaning abstracts away the details of any particular representation. Another possibility is that text could link to one or a subset of example stimuli (e.g. Fig. 4B, “exemplar”.) In such a case, the aspect of “meaning” would constitute representative visual examples of the word. The third possibility is that text links to all available visual representations (e.g. Fig. 4B, “available”.) If the visual representation reflects the consolidation of one’s experiential history with observed actions (*5, 33*) as expected for the consolidation of semantic memory, then neural responses to text may be understood as the activation of this consolidated visual experience. This suggests that a word’s meaning is uniquely rooted in an individuals’ experience.

The comparison between the predictions for these 3 models and the data strongly favor the available model. This architecture is also the easiest to implement, as a simple Hebbian mechanism will suffice and would predict that acquisition of verb meaning depends on the frequency of exposure, which has indeed been observed for several languages (*34*). The text response links only to a subset of the full distribution of visually selective units (Fig 4D). The reason for this is unclear but may reflect inefficiencies in the neural process that links verbs with visual representations and may be influenced by exposure or experience. In any case, reading a word does not evoke the same perceptual experience as viewing an action and thus substantial differences at the level of neural responses should be expected.

#### Fourth: origin of the link

Our results indicate that the link between text and the visual formats is not the product of imagery or learned associations that emerge from the task. We found that experimental manipulation of imagery has a minimal impact on the link between text and the visual formats. Further, semantic processing should be fast and automatic, and we found that it is exactly the early component of the correlation that was unaffected by the imagery manipulation (Fig 8D). In contrast, the late components of the correlation systematically differentiated passive viewing and imagery sessions (Fig 6C-E), demonstrating that the patient followed the instructions, a view also supported by the control experiment. The correlation between text and the visual formats was stable across all testing sessions (Fig 5). In our control experiment (Fig S7), passively viewing abstract symbols that had been paired to movement imagery did not induce a neural response. Thus our results are unlikely the consequence of learned associations. Yet another possibility is that neural responses to action verbs and observed actions represent implicit automatic motor plans (*35*). Indeed, how we plan and execute an action is an important component of meaning. However, our control study (Fig. S7) revealed no selectivity to passive observation of movement-predictive cues.

Another possibility is that the linked selectivity patterns for observed actions and action verbs reflect a population of cells that are responsive to the internal act of silent naming (i.e. generating the action verb). In this view, when viewing text or videos, the participant covertly generates the same word, and thus produces the same activity patterns. We think this hypothesis is unlikely. This hypothesis would predict results similar to the invariant hypothesis, as generating the action verb should be consistent across the different visual presentations. Instead, for simultaneously recorded neural populations, we find that text responses link to the visual formats in idiosyncratic ways (e.g. text and the lateral views, but not the frontal, Fig 1A, 4C). It remains possible that neurons are selective to particular cue-naming pairs, however, in our prior work (*36*) in the same participants, we found no selectivity for specific cue-intention pairings (e.g. response for imagined movement to the right when cued with a spatial target, but not when cued with symbol.) Thus, we think the naming hypothesis is unlikely.

### Nature of neuronal representation of variables coded in PPC

The ability of small neuronal populations to encode many variables is consistent with the mixed-selectivity scheme in which distributed, non-linear, high-dimensional representations code in a contextually dependent manner (*37, 38*). However, at least within the cortical locations explored in the current study, we find that such encoding is not random, but systematically organized around stimulus properties, a scheme referred to as partially-mixed selectivity (*39*). Indeed, neural populations coding the same basic-level action type for different formats overlapped (e.g. Fig 3, Fig S6). Partial mixed selectivity may represent a general structure for representing sensorimotor aspects of meaning within association cortices, resulting in rich links between text and the diversity of overlapping and distinct components of the visual formats that mirror the statistics of visual encoding independent of text (Fig 4). The partially-mixed architecture may also account for the weak link between text and the visual formats (e.g. relatively low population correlation and few units with matching SPs). If a cortical region encodes the many visual facets of an observed action (e.g. viewpoint, posture, and other untested features) and text links with both what is overlapping and distinct about action presentations, it follows that the link between text and any particular presentation must be relatively weak.

### Cortical organization of conceptual knowledge

In understanding an action verb, we access semantic knowledge. The cortical organization of semantic knowledge has been contentious. Some theories contend that conceptual knowledge is rooted in cortical regions that employ supramodal symbolic processing (*11*) while other theories take the opposite perspective, that semantic knowledge is encoded in the distributed sensorimotor network (*6, 40*). Most recent theories posit that meaning emerges from interactions between supramodal associative areas and regions directly responsible for processing sensory stimuli, motor actions, valence, and internal state (*1–5*); Our results are consistent with these interactions models, given the longer latencies we observed for text selective responses in PPC, relative to those reported for higher order language regions such as superior temporal gyrus, or inferior frontal gyrus (*41*). This model comes in many versions, primarily distinguished by which areas constitute the supramodal regions and the nature of the interactions. In part, a deeper understanding of the organization of conceptual knowledge in the human brain has been limited by the general inability to record from single neurons in humans. We know of no single unit recordings in supramodal regions, but one intriguing possibility is that these areas may host neurons similar to the “concept cells” of the medial temporal lobe (MTL, *42*), which respond to a preferred stimulus (e.g. a particular individual) largely independently of sensory modality or presentation details (e.g. image, written word, sound). While such strong invariance provides a model for neural coding mechanisms in supramodal centers, much less is known about how semantically related items are encoded in the distributed network. The current study contributes to this goal by providing the first demonstration of a link between words and their sensorimotor representations and how the neural architecture supports this link.

In the current paper, we have focused on how verbs are given meaning. We may also consider what our results mean from the reverse direction, how the neural population may contribute to naming an observed action. We find relatively high generalization across different views of the same observed action, but also a degree of dependence on viewpoint and the point of fixation (Fig S2). These neural properties suggest that rostral PPC neurons could play a role in creating increasingly abstracted representations that associate the same actions and thus contribute to the processing needed for naming, but, given the weakness of the link, that subsequent regions, potentially employing winner take all like mechanisms would be needed for the final conversion to labeling the observed action.

The link between visual representations of actions and action verbs fits with current views of how infants learn action verbs by mapping words onto conceptualizations of events (*43*). Infants can distinguish action exemplars (running, marching, jumping) independently of the actors (*44*) and that this ability predicts the use of action verbs at two years of age (*45*). Furthermore it provides an explanation for why infants learn verbs later than nouns (*46*), as the corresponding visual representations are in different visual pathways. In the PPC the development of observed action selectivity, which is originally in the service of guiding future actions (*22*), may only occur once the infant starts moving. Indeed infants initially learn verbs corresponding to their own actions (*47*).

### Limitations of the study

#### Stimuli

We used a restricted set of observed actions and action verbs, based on the category of actions that best evoke responses in neuroimaging (*48*). Recordings from the same arrays in the same subjects have demonstrated selectivity for imagined reaches and saccades (*36*), simple shoulder shrugs and vocalizations (*39*), as well as memory-strength and confidence (*49*). These diverse results point to additional functions that may also be recruited when processing the meaning of language that will require future investigations.

#### Visual Formats

We tested only a small number of visual formats: two postures and two viewpoints. Thus the visual invariance that we established may be an overestimation, and increasing the diversity of different presentations of the same action would lower the percentage of invariant cells. Hence, while it remains possible that the visual invariant neurons (F=L0=L1 in fig 4) are akin to concept cells as described in the MTL of humans, this is by no means established. To this point, neurons exhibiting invariance in the MTL showed sparse coding (only active for a single basic level category) while the invariant neurons tested in our study were broadly tuned, matching the tuning profiles of other visually selective neurons (Fig S4). The small number of visual formats may also partially account for text selective units with mismatched or absent visual selectivity (Fig. 4D, E) as they may link with other untested visual representations of the corresponding action identity.

#### Recording site

We tested only one region of the action observation network. Other regions of the action observation network (AON, e.g. premotor areas, or the LOTC), based on neuroimaging and lesion, likely play a role in linking language with its sensory and motor representations. Action verbs may be associated with the kinematic profiles of movement, movement dynamics, the agents typically performing the action, the objects typically subjected to the actions, the desired outcome or value of the action, the expected sensations that accompany the action, amongst others. The constituent regions of the AON likely encode these movement attributes and together may form the distributed network that links action verbs with these varied aspects of meaning.

#### Causality

As with all passive neural recording studies, our study cannot determine the causal role of our PPC neurons in understanding the meaning of action verbs. However, prior work has shown that damage or inactivation within the fronto-parietal AON can result in specific action comprehension deficits (*24, 27–29, 32*) consistent with the idea that neurons within the AON play an important role in verb comprehension. Our results provide clarity on the presence and nature of the link between neural representations of action verbs and visually observed actions at the level of single units in PPC.

## Conclusion

The current study provides the first single unit evidence that action verbs share a neural substrate with visually observed actions in high-level sensory-motor cortex thus clarifying the neural organization of human conceptual knowledge. Action verbs link with all the diverse visual representations of the related concept suggesting that language may activate the consolidated visual experience of the reader.

## Materials and methods

### Experimental Design

#### Data acquisition

All procedures were approved by the California Institute of Technology, University of California, Los Angeles, and Casa Colina Hospital and Centers for Healthcare Institutional Review Boards. Informed consent was obtained from NS & EGS after the nature of the study and possible risks were explained. Study sessions occurred at Casa Colina Hospital and Centers for Healthcare and Rancho Los Amigos National Rehabilitation Center.

#### Behavioral Setup

All tasks were performed with NS & EGS seated in their motorized wheel chair. Tasks were displayed on a 28 inch or 47-inch LCD monitor. The monitors were positioned so that the screen occupied approximately 25 degrees of visual angle. Stimulus presentation was controlled using the Psychophysics Toolbox for MATLAB.

#### Physiological Recordings

NS and EGS were implanted, each with one 96-channel Neuroport array on the gyrus dorsal to the junction of the intraparietal (IPS) and post-central (PCS) sulci (Fig S1). These locations were implanted based on three considerations: first, the Neuroport arrays used in the current study are not suitable for implantation within sulci given short electrode shank lengths (<= 1.5mm) and lack of long-term viability for direct implantation within sulci. Thus, implant locations must be restricted to gyri accessible on the cortical surface. Second, the cortical regions of interest are near the junction of the IPS and PCS. This consideration was included as we were targeting functional responses related to grasping, manipulation, and other behaviors that emphasize the hand. Cortical regions within and around the junction of the IPS and PCS in human neuroimaging studies have consistently shown preferential responses to hand-based actions. Third, we used functional magnetic resonance imaging within the individual participants to identify regions with a preferential response for grasping actions. We used two neuroimaging tasks suitable for paralyzed individuals, to identify grasp related responses in each individual subject. The resulting functional responses, combined with the constraints described above, determined the implant locations are shown in supplementary figure 1.

Grasp related responses around the junction of the PCS and IPS have typically been described as the anterior IPS or putative human homologue of the anterior intraparietal area (phAIP) and is generally assumed to be the human homologue of macaque AIP. Macaque AIP is a region localized to the *lateral* bank of the anterior portion of the intraparietal sulcus involved in the visual control of grasping actions. However, the *medial* bank of the anterior intraparietal sulcus contains a distinct grasp field. This grasp field, described as PEip or Brodmann’s area 5L (BA5L), is characterized by distinct frontoparietal connections (AIP is densely interconnected with PMv while PEip is connected with the rostral portion of M1), direct connections to the hand regions of the spinal cord, bilateral somatosensory responses, and functional responses related to hand and finger movements. While some progress has been made in the identification of the human homologue of AIP (2), the human homologue of PEip/BA5L has not yet been established, and it may be the case that neuroimaging results around the junction of the PCS and IPS include both the human homologues of macaque AIP and PEip/BA5L. Additional work probing the single unit properties of the arrays in the two human subjects is needed to better understand the functional homologies of the regions investigated in the current study. In light of this uncertainty we refer to the recording sites as parietal grasping regions.

Neural activity was amplified, digitized, and recorded at 30KHz with the Neuroport neural signal processor (NSP). The Neuroport System, comprising the arrays and NSP, has received FDA clearance for <30 days acute recordings; for purposes of this study we received FDA IDE clearance (IDE #G120096,) for extending the duration of the implant.

Single and multiunit activity was sorted using k-mediods clustering using the gap criteria to determine the total number of neural clusters. Clustering was performed on the first n principal components where n was selected to account for 95% of waveform variance (range of 2-4 components.) Sorting was reviewed and adjusted if deemed necessary following standard practice by merging or splitting clusters as needed.

#### Tasks and Stimuli

Experimental stimuli consisted of five manipulative actions (drag, drop, grasp, push, rotate) displayed in four different “formats”: three visual video formats and one text format. In two visual formats the actors were viewed from the side but the actor was either standing next to a table (Lateral 0, L0) or sitting in a lotus position on the floor (Lateral 1, L1). In the third visual format, the actor was standing next to the table, but was viewed from the front. Thus L1 differed from L0 only by body posture; F from L0 only by viewpoint (note that two video cameras were used to simultaneously acquired videos for F and L0 and thus the timing and kinematics of the movements are identical), and F from L1 both by viewpoint and posture. In the text condition, the written action word (Arial at font size 80) was shown for 2.6s. Experimental stimuli of the visual formats consisted of video-clips (448×336 pixels, 50 frames/s) showing one actor at a distance of 1.2 m performing five different hand actions (drag, drop, grasp, push, rotate) directed towards an object (four versions per action-format). The objects were positioned directly adjacent or within the hand such that actions predominately involved the wrist and fingers. All videos measured 17.7° by 13.2° and lasted 2.6 s (the first 2 and the last 2 frames being static). The edges of the videos were blurred with an elliptical mask (14.3°×9.6°), leaving the actor and the background of the video unchanged, but blending it gradually and smoothly into the background around the edges. We did not enforce fixation, instead asking the subject to view the actions in a naturalistic manner. The effects of fixation were documented in a separate set of experimental sessions described below. Each data session consisted of 12 repetitions of each unique video. Presentation was split into 3 runs (4 repetitions per run, corresponding to the four versions). Videos were presented in a pseudorandom manner: all conditions were randomly ordered and presented once before repetition. During baseline a highly blurred (full-width half-max = 80 pixels) static frame (average of all video frames) was shown.

We collected this data under two different instruction sets. In the first instruction set the subject was instructed to attend but otherwise passively view experimental stimuli. For the text format, we asked the subject to read the word silently without any accompanying visualization. We collected 7 (subject NS) and 5 (subject EGS) sessions in this passive viewing paradigm. In the second instruction set, the subject NS (6 sessions) was instructed to use the text as a prompt to visualize the associated action being performed from either the frontal (F, 3 sessions) or lateral standing perspective (L0, 3 sessions). Six datasets in total were collected over 3 sessions. Each dataset was collected in full under the same instruction. The datasets were collected over 3 experimental sessions in the following order (F, L0), (L0, F), (F, L0) with () indicating that the datasets were acquired during the same session. Prior to each run, the subject was reminded of the experimental condition. Following each run, the subject reported which perspective she used to imagine the actions.

Information on the preliminary action observation task (Fig S2) to test for the presence of units selective to observed actions is described in supplemental information. Additional information is found in supplemental materials.

### Statistical Analysis

#### Within-format individual neuron analyses

##### Individual unit event related averages (Fig 1A)

For each unit, neural activity was averaged within a 750 ms window starting from −1.5s before video onset and stepping to 4 seconds with 100 ms step intervals. Neural responses were grouped by the observed action exemplar and a mean and standard error of the mean (n=12) were computed for each time window and for each action.

##### Within-format one-way ANOVA for action identity (Fig. 1B)

We defined a unit to be selective for action identity for a particular format if the unit displayed a significant differential firing rate for the five action types during video presentation. Firing rate was taken as the total spike count during movie presentation (2.6 seconds) starting at 0.25 seconds after onset divided by the window duration. Firing rates were subjected to a one-way ANOVA with the factor of action identity and significance was determined as p<0.05 after fdr correction. To ensure that selectivity was driven by the task stimuli, we repeated the analyses using the 1 second window prior to stimulus onset and found no neurons (0%) were selective in any format after fdr correction.

##### Within format linear fit of neural responses (Percent selective, cross-validated coefficient of determination R^2^, and coverage)

We used a linear regression model for multiple analyses. For each neuron, we fit a linear regression model that explained neural firing rate during movie presentation as a function of the categorical variable manipulative action identity (five actions: drag, drop, grasp, push, rotate). For some analyses, linear fits were performed separately for each of the four formats (e.g. written verb, frontal view, and the two lateral views). For others, such as the model selection procedure described below, we included various combinations of formats with constraints on the beta parameters (described below). Firing rate was taken as the average unit response in a single window that extended for the duration of movie presentation (2.6 seconds) starting at 0.25 seconds after onset. The baseline response was taken as the average activity in the 1 second preceding movie onset.

###### Coverage (Fig S3)

We computed the p-value associated with the coefficient estimate of each action (e.g. drag, drop, etc.) and found the percentage of coefficients associated with each action having a p-value less than 0.05 after fdr correction. We used a bootstrap procedure for generating 95% confidence bounds on the estimates of the percent selective.

We also wanted to know the frequency with which the different actions resulted in the peak response. To test this, we first split the data into excitatory and inhibitory units. Inhibitory units were defined as units for which the beta coefficient for all five actions was negative and thus suppressed relative to the baseline response. We then identified the action that resulted in the largest deviation from baseline activity and counted the number of units that showed a peak response for each action.

###### Cross-validated coefficient of determination (cvR^2^) (Fig 1C)

To derive a measure of strength of selectivity, we performed a leave-one-out cross-validation procedure to estimate the cross-validated coefficient of determination (cvR^2^.) For each fold, the single unit regression was parameterized using all but one trial, and the resulting model was used to predict the firing rate of the held-out trial. This was repeated and the R^2^ was computed using the held-out data as 1− (sum of squares of residuals)/(total sum of squares). The 95% confidence interval was estimated with a bootstrap procedure.

##### Selectivity curve analyses (Fig S4)

Is neural coding for observed actions sharp or graded? For each selective unit (within-format ANOVA p<0.05 fdr corrected, see above) within each format, repetitions for each observed action were split in half to create training and testing splits of the data. Repetitions were then averaged to create a single value per action for each of the training and test sets. Training set data was rank ordered from the action resulting in the highest firing rate to the action resulting in the lowest firing rate. This computed order was used to sort the test data. This process was repeated 500 times and the results were averaged across folds. The result is a cross-validated measure of the response of each unit as a function of rank. Responses were normalized between 0 (response to worst action computed from the *training data*) and 1 (response to best action computed from the *training data*) before averaging across the population of selective units. Confidence intervals were estimated using a bootstrap procedure. Both the mean with 95% confidence interval (estimated with a bootstrap procedure) and the full distribution (shown in a violin plot) are presented.

#### Within-format neuron population analyses

##### Time resolved classification of action exemplars (Fig 1D)

We performed a sliding window classification analyses to measure the strength and latency of population coding of observed actions and action verbs. For each time window, we constructed a classifier to differentiate the observed action for each format separately. Classification analyses was performed using linear discriminate analysis with the following assumptions: one, the prior probability across the action exemplars was uniform, two, the conditional probability distribution of each unit on any given action exemplar was normal, three, only the mean firing rates differ for each action exemplar (the covariance of the normal distributions were the same for each action exemplar), and, four, the firing rates of each input are independent (covariance of the normal distribution was diagonal). Relaxing these constraints (e.g. allowing a full-rank covariance matrix) generally resulted in poorer generalization performance. The classifier took as input a matrix of average firing rates for each sorted unit. We did not limit analyses to action selective units to avoid “peeking” effects. Classification performance is reported as generalization accuracy of a stratified leave-one-out cross-validation analysis. The average neural response was calculated within 300 ms windows, stepped at 10 ms intervals. Window onsets started from −.75 seconds relative to video onset with the final window chosen to be +3 seconds. The window size was chosen to ensure that a reasonable estimate of firing rate could be determined while still allowing temporal localization. Classification was performed on all sorted units acquired within a single session. Mean and bootstrapped 95% confidence intervals were computed for each time bin from the cross-validated accuracy values computed for each session.

###### Latency analyses (Fig 1D)

We used a method to identify the abrupt change in classification accuracy as the signal transitioned from chance decoding levels prior to stimulus onset to significant decoding. For each session, accuracy as a function of time was split into two regions (an early and late period) and the sum of the squared residuals (SSR) was computed after subtracting the mean. The time of transition between the early and late period was stepped through each time bin defined in the sliding window classification analyses. Latency is reported as the time of the abrupt change as defined by the transition time that resulted in the lowest SSR. This procedure resulted in 13 latency estimates for each format, (one for each of the 13 sessions in subject NS.) We used a 1×4 ANOVA to test for a main effect of format on latency followed by post-hoc testing between pairs of formats. Latency analyses was not attempted for subject EGS as decode accuracy was worse and we had fewer sessions (5 sessions for EGS versus 13 sessions for NS) although the general time-course of decode accuracy was similar (see Fig 2C).

#### Cross format individual neuron analyses

##### Comparing selectivity patterns between formats. (Fig 3, Fig 4, Fig S6)

How a unit codes for action identity within a format is described by its selectivity pattern (SP) defined as the precise firing rate values for all five action identities (see Fig 3A). At the single unit level, understanding how a unit codes action idenity across formats can be quantified by comparing SPs across formats. There are four possibilities when comparing SPs across a pair of formats: **one,** the SPs are similar or matched across formats, **two,** both formats are selective but the SPs are distinct across formats, **three**, only the first of the two formats is selective for action identity, and, **four**, only the second format is selective for action identity. Each of these possibilities can be defined mathematically within a linear models framework using four models of neural coding across formats:

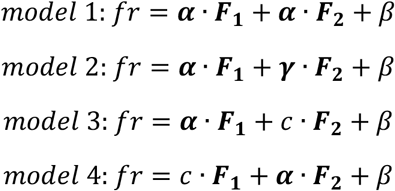

In model 1, the linear fit is constructed with the constraint that the weight parameters (***α**ϵR*^5^) for each action exemplar are the same across the two tested formats (*F*_1_, *F*_2_). This model describes units with the same SP across formats. In model 2, the weight parameters (***α, γ**ϵR*^5^) are allowed to be different, and enable distinct selectivity patterns for action exemplars between the two formats. In models 3&4, one format is assumed to be unmodulated by action identity and a single scalar value (*cϵR*^1^) describes the presumably equivalent response (e.g. non-selective) to all actions within the format.

To determine how SPs compared across formats we fit the parameters of each of the four models using standard linear regression techniques (see above) and the results were compared. Several measures are commonly used to select the “best” model from a set of candidate models. We used both the Bayesian information criterion (BIC) and cross-validated R^2^ (cvR^2^) as our model selection criteria. These two methods provided slightly differing but reasonable notions of similarity. Heuristically, cvR^2^ required a near exact match while BIC was found to have a more qualitative notion of similarity (See Fig S6). We viewed the two measures as providing something akin to upper and lower bounds on whether units were similar across formats, and do not view either method as being “correct” per se. We used the arithmetic mean of the percentages of units fitting each model when reporting the results in the main figures of the paper.

This analysis was extended to 3 formats (fig 4A) requiring the creation and evaluation of 15 models for all the unique ways that the SPs can be expressed across formats. The 15 models are enumerated below (Models 1-15) as the set of coefficients for each format where, following the description for pairwise comparisons, equivalent Greek letters indicate matched selectivity patterns across formats, a constant *c* indicates no selectivity to action exemplar for the associated format, and multiple Greek letters indicates significant but idiosyncratic selectivity patterns across formats. We performed this analyses for all combinations of three formats. Finally, the analysis was extended to 4 formats requiring the creation and evaluation of 51 models for all the unique ways that the SPs can be expressed across formats. These models were constructed by enumerating all the possible configurations that the “fourth” format can take relative to the 15 models described above. This analyses pooled data from subject NS and EGS given the relative paucity of data for EGS. The 15 possibilities (3 formats) are shown below:

| Model # | Format 1 | Format 2 | Format 3 | Format 4 (possible values) |
| --- | --- | --- | --- | --- |
| 1 | $\alpha$ | $\alpha$ | $\alpha$ | $(\alpha, \gamma, c)$ |
| 2 | $\alpha$ | $\alpha$ | $\gamma$ | $(\alpha, \gamma, \beta, c)$ |
| 3 | $\alpha$ | $\alpha$ | $c$ | $(\alpha, \gamma, c)$ |
| 4 | $\gamma$ | $\alpha$ | $\alpha$ | $(\alpha, \gamma, \beta, c)$ |
| 5 | $c$ | $\alpha$ | $\alpha$ | $(\alpha, \gamma, c)$ |
| 6 | $\alpha$ | $\gamma$ | $\alpha$ | $(\alpha, \gamma, \beta, c)$ |
| 7 | $\alpha$ | $c$ | $\alpha$ | $(\alpha, \gamma, c)$ |
| 6 | $\alpha$ | $c$ | $c$ | $(\alpha, \gamma, c)$ |
| 9 | $c$ | $\alpha$ | $c$ | $(\alpha, \gamma, c)$ |
| 10 | $c$ | $c$ | $\alpha$ | $(\alpha, \gamma, c)$ |
| 11 | $\alpha$ | $\gamma$ | $c$ | $(\alpha, \gamma, \beta, c)$ |
| 12 | $c$ | $\alpha$ | $\gamma$ | $(\alpha, \gamma, \beta, c)$ |
| 13 | $\alpha$ | $c$ | $\gamma$ | $(\alpha, \gamma, \beta, c)$ |
| 14 | $\alpha$ | $\gamma$ | $\beta$ | $(\alpha, \gamma, \beta, c)$ |
| 15 | $c$ | $c$ | $c$ | $(\alpha, c)$ |

For 4C we explicitly restricted the distribution of cells to those that matched specific selectivity patterns between text and the visual formats.

#### Cross format neuronal population analyses

##### Cross-decoding analyses (Fig 2)

We used two methods to measure the similarity of population responses across formats: cross-format decode accuracy and population correlation.

Classification analyses was performed using linear discriminate analysis with assumptions and cross-validation procedures as described for within-format decoding above. For cross-format results, classifiers were trained within format and applied to alternate formats. More precisely, for each fold of the within-format cross-validation procedure, the classifier was applied to the neural data associated with each of the three other formats. All predictions across folds of the cross-validation procedure were used to compute decode accuracy. This enables us to understand how well the neural representation of the different action exemplars generalize to a novel format when the definitions of the actions are preserved across the two formats. This approach further introduces directionality to the comparisons: e.g. how well do definitions established for the text format generalize to the visual formats and vice versa. To verify that the ability to generalize from one format to another required correctly aligning the action exemplars across formats, we repeated the analyses but now using “mismatched” labels. In the mismatch analyses, the action identity labels were swapped between action exemplars and accuracy was recomputed based on these reassigned labels. For the mismatched condition, we performed all possible shuffles for which no action exemplar was matched across formats (N=44).

##### Cross-format *population* correlation analyses (Fig 2, Fig 5, Fig 6)

To compute the population correlation measure we organized the neural response data into 4 vectors, one for each format (Fig S5.) Each vector had five values per unit, one value for each of the five actions. This value was computed as the mean firing rate recorded during the 2.6 seconds of stimulus presentation offset by 0.25 ms, averaged across trial repetitions. The mean response across these five values was subtracted and the five values per unit were then concatenated across units to create a population response vector for each format. The same procedure was performed for each format ensuring that the same units and actions were aligned across the format vectors. The Pearson correlation was computed across format vectors to quantify the population level similarity. Note that subtraction of the mean response across the five actions prior to concatenation was done to ensure that a positive correlation value across formats reflected similarity in the pattern of responses to the five actions and not general offsets in the mean response of the different neurons. This was necessary as some units were activated above baseline for all actions and some were inhibited below baseline for all actions biasing the population towards a positive correlation that was not driven by the patterns of selectivity for the different actions. To ensure that a significant correlation was specifically the product of comparing the responses to the same actions across formats we performed a shuffle control analysis. The population correlation was computed using the same procedures except that the five values computed per unit, one for each action identity, were misaligned (shuffled) between formats. The same shuffle order was applied to all units. All possible ways of shuffling action identities between formats (e.g. reordering 5 values) were tested and the resulting shuffled correlations were averaged in reporting the results. The fiducial and shuffled correlations were computed separately for each session. Significant population correlation was determined based on the p-value resulting from a one-sided t-test to determine whether the distribution of correlation values computed for each session was greater than 0. The correlation values for each session are also shown separately for each session with 95% confidence intervals computed using a bootstrap procedure (see figure 5).

The correlation analysis was also performed using a sliding window approach to look at the time scale of positive correlation between formats (fig 5). The approach was similar as described with the following modifications: (1) Because we computed within-format correlation in addition to across-format correlations, we used a cross-validation approach for computing the correlation values. The same procedure as described above was performed, however, the process began with splitting trial repetitions into training and testing sets (6 trial repetitions each) and concluded with computing the correlation across the training and test splits. (2) The training and test sets were computed from windowed data. Windows centered on time x were computed using a pseudo-Gaussian weighting function with mean=x, and standard deviation – 200ms. This allowed for a relatively smooth and temporally precise measure of the neural response. Correlations were computed between training and test sets for all combinations of windows starting from 500ms before movie onset to 500ms after movie offset.

We further asked whether the correlation between any two formats was mediated by the remaining two formats. For instance, the correlation between the text response and the frontal view could be the consequence of a text being correlated with the lateral view and the lateral view being correlated with the frontal view. To address this possibility, we performed a partial correlation analyses with the neural data from all four formats, thus looking at the correlation between two formats while regressing out the shared variance with the remaining formats.

#### Control analyses and tests

##### Understanding the effect of explicit visual imagery (Fig 6)

Can the text response be understood as the consequence of visual imagery: replaying the visual stimuli by imaging the visual sequence of events? In analyzing the visual formats we found that the frontal and lateral view were encoded in a distinct manner. For instance, only roughly half the units had a matched selectivity pattern across the frontal and lateral perspectives while the remaining population of selective units had a distinct selectivity pattern. If the neural responses depend on the contents of visual imagery, then visualizing from the frontal perspective should tend to activate the selectivity patterns for the frontal view while visualizing from the lateral perspective should tend to activate the selectivity patterns from the lateral perspective. To understand the impact of visual imagery we split the data based on the task instructions given to the subject prior to each session. In comparing the results of the model selection analysis, we compared the percentages of cells classified into each category (e.g. the percentage of cells with matched selectivity across all formats, a single format, etc.) using a chi-squared test.

In addition, we split the dynamic correlation results into sessions in which the participant was instructed to passively view (7 sessions) or actively imagine movements (6 sessions) when presented with the action verb. We performed a paired t-test between these two groupings at each pixel location to test for significant changes in correlation value as a function of task instruction (significance tested at the p<0.05 level). The correlation values at the significant pixels were averaged and plotted to visualize the shape of temporal trends in correlation values.

##### Supplemental control task (Fig S7)

We performed a sensory-motor association learning task to test whether repeated presentation of abstract stimuli, when paired with motor imagery of an action, would result in neural selectivity under passive viewing conditions. In the context of the current paper this helps to constrain the interpretation of a shared neural substrate for action verbs and visually observed actions. We instructed the subject to use visual imagery to imagine finger movements when presented with fractal-like images of snowflakes. We used five images. Three of the images were associated with visual imagery of finger flexion movements of the thumb, index, and ring fingers. These movements were chosen as they resulted in especially robust neural selectivity in preliminary testing. Two additional images were used as controls; the subject was instructed to passively view the stimuli without accompanying visualization. These two control images were used to test whether differential responses might emerge between the two images based on repeated exposure, even in the absence of any overt behavior on the part of the subject. We found that no such differential tuning emerged and thus, for the current study, only one of the control images was used in the analysis.

The experiment began with a passive viewing session in which the subject viewed the stimuli prior to any motor association to test for base-line visual selectivity. Then, for the first 16 repetitions of each condition during the first two session days, the snowflakes were presented along with a key instructing which action should be performed (or no action in the case of the control images). For all other trial repetitions, the key was removed and the subject performed reaction time and delayed imagined movements when presented with the visual stimuli. At varied intervals (Fig 8), we asked the subject to passively view the same set of stimuli. The experiment was performed twice, sequentially, with each experiment similar in structure but using a different set of visual stimuli. Experiment 1 consisted of 328 total repetitions per stimulus presented over 14 session days in a 49 day period. Experiment 2 consisted of 264 total repetitions per stimulus presented over 10 session days in a 56 day period.

We used a time-resolved classification analysis on the passive and reaction time trials separately to quantify selectivity for the cued stimuli. Classification methods were the same as described above (Time resolved classification of action type) with windows beginning at −.5 seconds and stepping to 2 seconds. For decode accuracy of the passive stimuli as a function of session number, we used the average firing rate within a 1 second window offset by 250 ms from stimulus onset.

## Supplemental materials

Figure S1: Functional magnetic resonance imaging (fMRI) localization of implant sites.

Figure S2: Preliminary study; impact of multiple factors on neural responses to observed manipulative actions.

Figure S3: Percent of selective units for each action for each of the four formats.

Figure S4: Shape of tuning functions for each format.

Figure S5: Organization of data for population correlation method.

Figure S6: Comparison of model selection procedures.

Figure S7: Abstract symbols systematically associated with motor imagery does not evoke differential neural tuning.

## Acknowledgments

This work was supported by the National Institute of Health (R01EY015545), the Tianqiao and Chrissy Chen Brain-machine Interface Center at Caltech, the Conte Center for Social Decision Making at Caltech (P50MH094258), the Boswell Foundation, and ERC (Parietal action) VII FP (323606). The authors would also like to thank subject N.S. and E.G.S. for participating in the studies, Viktor Scherbatyuk for technical assistance, and Kelsie Pejsa for administrative and regulatory assistance.

We would also like to thank Mick Rugg for helpful comments on an early version of this manuscript.

## Author contributions

Conceptualization, T.A.; Methodology, T.A. and G.A.O.; Investigation, T.A. and C.Z.; Formal Analysis T.A.; Writing – Original Draft, T.A.; Writing – Review & Editing, T.A., G.A.O., and R.A.A.; Funding Acquisition, T.A., G.A.O., and R.A.A.; Resources, E.R. and N.P.; Supervision, T.A., G.A.O., and R.A.A.

