## Supplemental Materials and Figures for "A shared neural substrate for action verbs and observed actions in human posterior parietal cortex"

### Supplementary Materials

**Preliminary action observation experiment (Fig S2)**

Experimental stimuli consisted of video-clips (448x336 pixels, 50 frames/s) showing an actor, sitting at a table, viewed from the side at a distance of 1.2 m, performing seven different hand actions directed towards objects. The action videos used for this preliminary experiment are the same as (*22*). Each of these seven actions (drag, drop, grasp, push, roll, rotate, squeeze) was performed by a male or female, using two natural objects (a plumb and a mandarin), yielding four versions of each action exemplar. The objects lay on a table or in the hand, with the arms on the table until the actions began, hence actions predominately involved the wrist and fingers. We took extensive precautions to equate videos across the actions: the same two actors, wearing the same clothing performed the actions without expressing any particular emotion. Efforts were also made to equalize the scenes in which the action took place. Lighting and background were identical as well as the general organization of the scene. The actor stood on the right, kept his/her hands in the middle when performing the action.

All videos measured 17.7° by 13.2° and lasted 2.6 s (the first 2 and the last 2 frames being static). Before the presentation of each video, a blurred static frame of the following video was shown as baseline. The edges of the videos were blurred with an elliptical mask (14.3°x9.6°), leaving the actor and the background of the video unchanged, but blending it gradually and smoothly into the background around the edges. A 0.2° fixation target was shown in all conditions. The fixation target was presented as close as possible to the position where the movement took place in the video, either above or below the midpoint of the trajectory of the manipulating hand. Each data session consisted 10 repetitions of each unique video for a total of 560 videos watched per session (2 actors x 2 objects x 2 fixation locations x 7 actions x 10 repetitions). Presentation was split into 3 runs (3 + 3 + 4 repetitions). Videos were presented in a pseudorandom manner: all conditions were randomly ordered and presented once before repetition. The subjects were instructed to attend to but otherwise passively view the videos.

**Multifactor sliding window ANOVA for the preliminary experiment.**

In the preliminary experiment we manipulated 4 experimental variables that included action exemplar, fixation location, gender, and object. To establish whether these variables impacted neural responses we performed a sliding window ANOVA analyses. For each time window, we constructed an ANOVA model to explain firing rate for each unit using all factors (action, gender, object, fixation position) and their interactions. The resulting p-value for each factor and each time window was fdr corrected to determine significance. Window onsets started from -1.5 seconds relative to video onset with the final window chosen to be +3 seconds. Windows were stepped at 100 ms intervals. Each window included the average neural response within a 750 ms window.

### Supplementary Figures

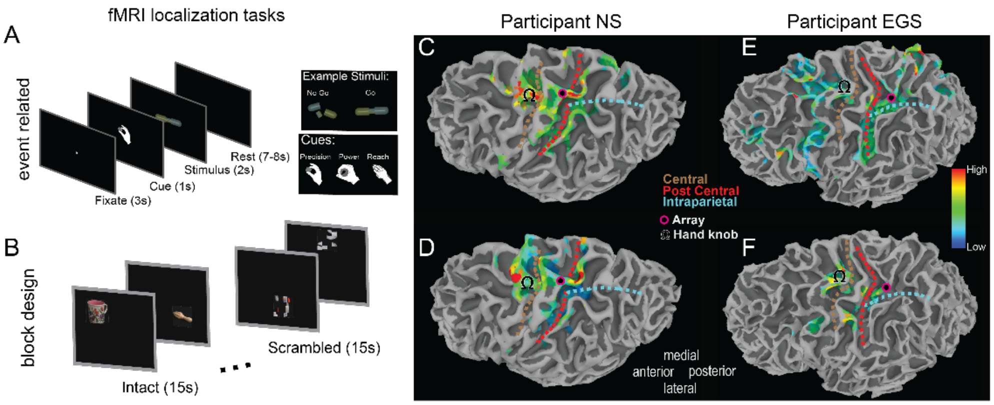


**Figure S1: Functional magnetic resonance imaging (fMRI) localization of implant sites.** fMRI was used to identify cortical regions that were suitable for implantation based on the BOLD response for imagined reaching and grasping actions. We performed complementary tasks to ensure robust activation across a range of paradigms and protocol types. **(A)** Event related task design. Following an intertrial interval, the subject was cued to perform a specific imagined movement (precision grasp, power grasp, or reach without hand shaping.) Following the cue, a cyndrilical object was presented. If the object was intact, the subject performed the cued imagined movement. If the object was broken, the subject withheld movement. **(B)** Block task design. Eight blocks were presented for 30 seconds per run. During the first 15 seconds common objects were presented with a new image presented every three seconds at a new spatial location. The subject was instructed to either imagine pointing at, imagine reaching and grasping, or look at the object in a natural manner prior to each run. During the last 15 seconds of each block, scrambled images were presented and the subject was instructed to guess the identity of the object. **(C)** Statistical parametric map showing voxels with significant activity in the grasp "Go" versus "No-Go" condition ($p<0.01$ fdr corrected.) Array location and cortical landmarks as depicted in the legend. **(D)** Statistical parametric map showing voxels with significant activation ($p<0.01$ fdr corrected) for the grasping versus looking condition in the block design. **(E,F)** Same as C, D but for participant EGS. See methods: "Data acquisition" for additional implant location details.

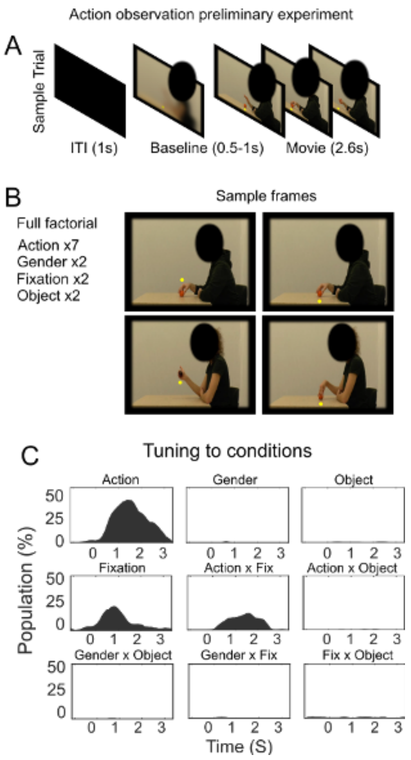


**Figure S2: Preliminary study; impact of multiple factors on neural responses to observed manipulative actions.** In a preliminary study we measured the impact of several conditions on neural responses from a single lateral perspective (similar to L0). **(A)** Sample trial illustrating temporal progression through first experiment. Face is masked to obscure identity per publisher's request. **(B)** Full factorial design used for testing neural selectivity for observed manipulative actions (OMAs). Each subject passively viewed pre-recorded videos of seven different manipulative action exemplars (dropping, dragging, pushing, rolling, rotating, grasping, and squeezing) while varying actor gender (male, female), object (orange, plum), and fixation location (above, below). Sample frames illustrate stimuli. Face is masked to obscure identity per publisher's request. **(C)** The percent of recorded units with significant responses to the factors of interest (ANOVA $p<0.05$, fdr corrected) through time. In addition to observed action, fixation location has a sizeable and significant impact on the neural population. Data represents the pooled population of units recorded from participant NS (3 sessions, 290 total units) and EGS (3 sessions, 111 total units).

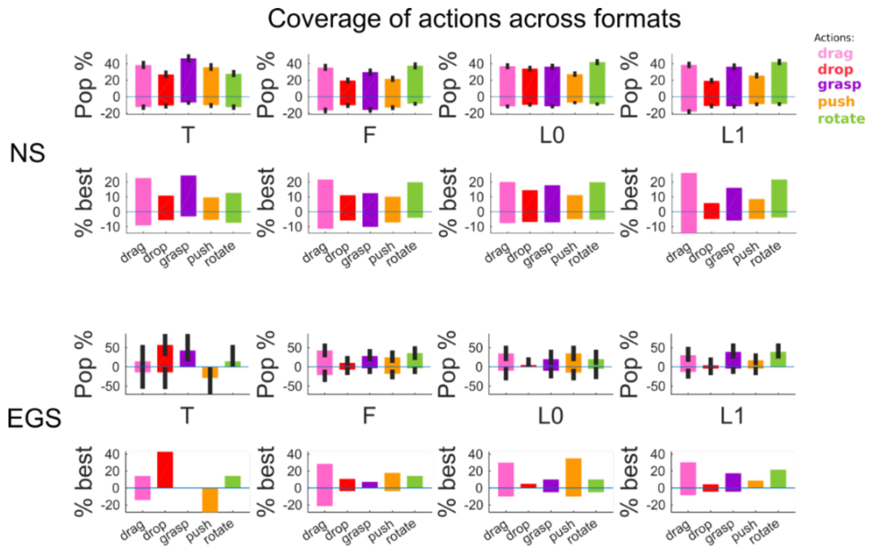


**Figure S3: Percent of selective units for each action for each of the four formats.** Results are shown individually for each subject. *Top*: Percent of units with significant modulation to each action type during presentation relative to baseline activity (paired t-test, $p<0.05$ fdr corrected, mean ± 95% CI) *Bottom*: Percent of units whose highest magnitude response belonged to each action type. Since roughly 1/3 of all recorded neurons showed purely suppressive responses (all actions suppressed activity below baseline), analysis is split between excitatory and purely suppressive units. Suppressive units shown as "negative" percentages. Color code for each action provided in inset. Formats shown in labeled columns (T=text, F = front, L0=lateral standing, L1 = lateral lotus).

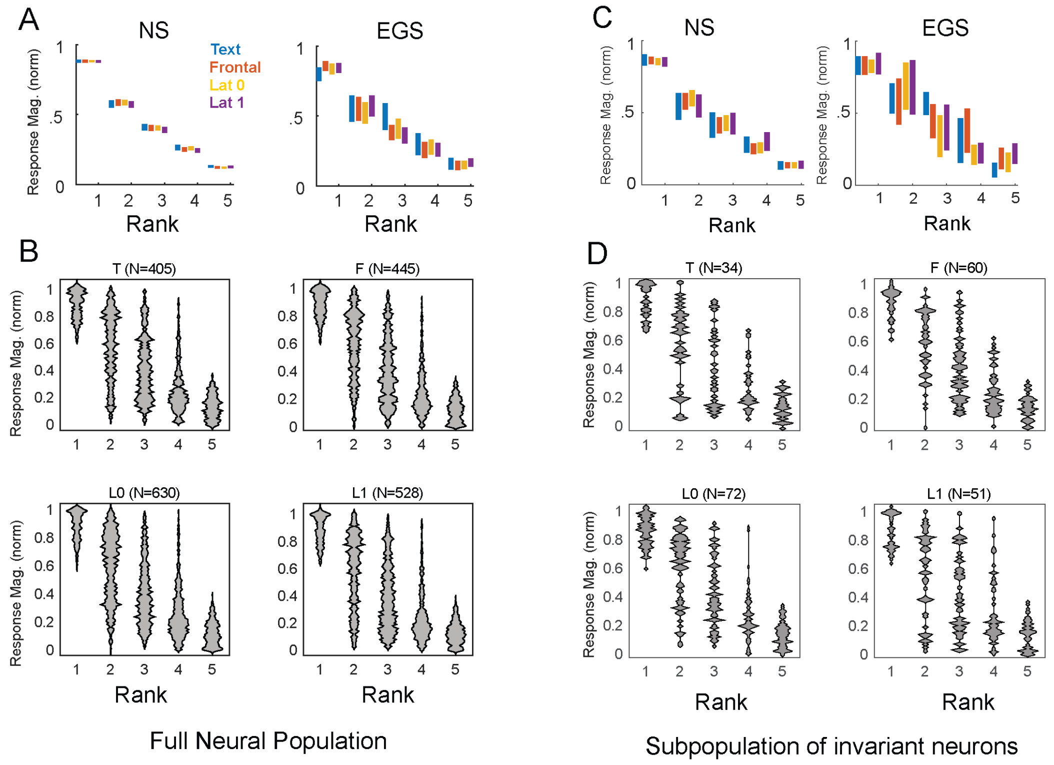


**Figure S4: Shape of tuning functions for each format. (A)** 95% confidence interval for normalized, cross-validated rank ordered responses for each of the four formats (see color code) and both subjects. Responses show a graded drop-off from the best action (defined as resulting in the largest magnitude response) second best action, etc. **(B)** Grey violin plot shows the probability distribution of cross-validated rank-ordered responses for subject NS. **(C)** Same as A, except for the subpopulation of cells that demonstrated invariance across all formats (invariance defined as T=F=L0=L1, Fig. 5B). **(D)** Same as B, except for the subpopulation of invariant neurons.

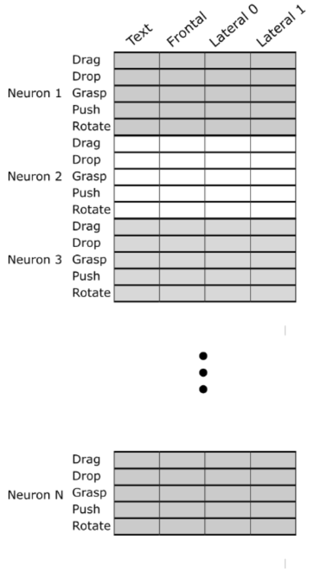


**Figure S5: Organization of data for population correlation method.** In order to compute a similarity measure between formats: the data was organized into matrix form such that the responses for the same unit and same action identity were positioned at corresponding rows across formats (as shown in the figure.) The correlation measure was computed across columns in order to provide a quantitative measure of how similar the pattern of responses for action identities was across formats.

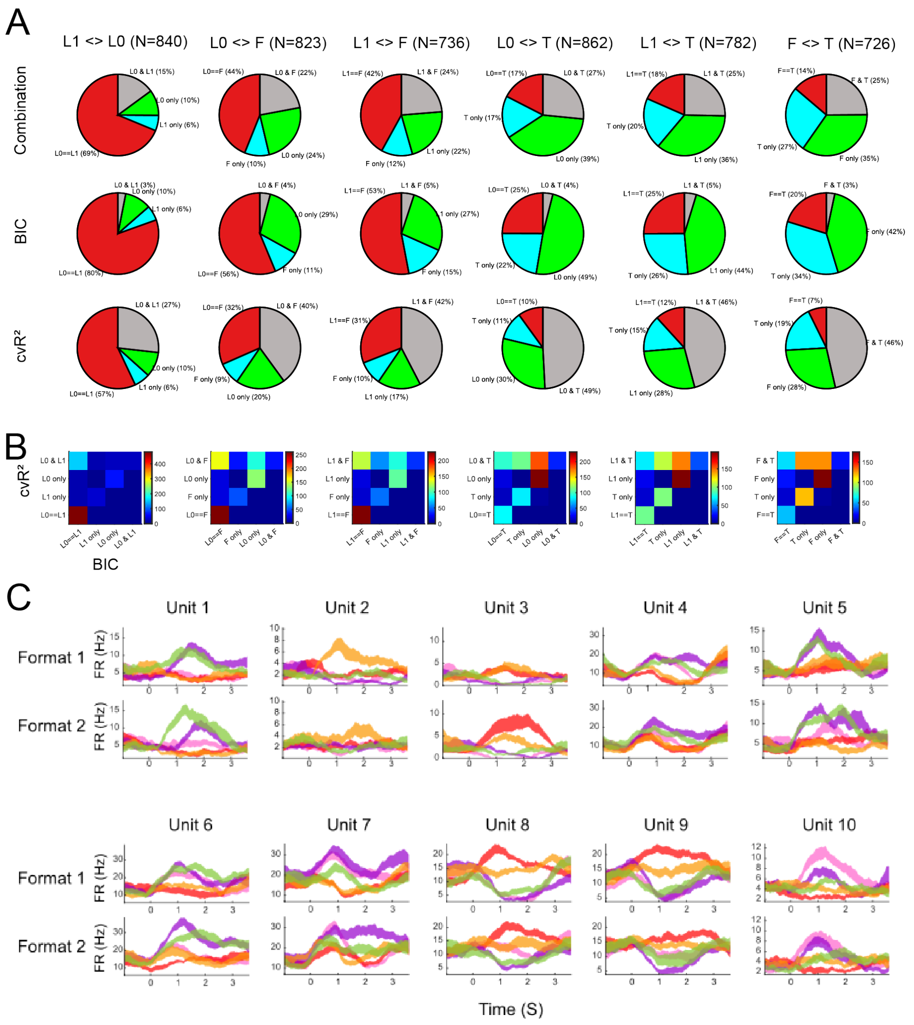

**Figure S6: Comparison of model selection procedures.** In main figure 3B we averaged the results of using Bayesian information criteria (BIC) and cvR2 as the criteria to identify the four (red, grey, green and cyan correspond to the four panels shown in 3A) selectivity patterns across formats. **(A)** The results are extended to include all possible pairwise comparisons of formats (top row: Combination) as well as results using Bayesian information criterion alone (middle row: BIC) and cross-validated coefficient of determination alone (bottom row: cvR$^2$). Results are organized by the comparison metric (rows) and the formats used in the comparison (columns). The combination measure is the arithmetic mean between BIC and cvR2. Only action selective units were used in the model selection analyses (Total number of selective units for the comparison shown as N=xxx in the title.) Selective units were defined as significant coding for action identity in a 2x5 format by action ANOVA with interactions (p-value <0.05 fdr corrected, either main effect or interaction.) The results are shown for subject NS. Results for EGS were similar. **(B)** A "confusion matrix" summarizing how units categorized using BIC were categorized using cvR$^2$ across the population. For instance, >400 units categorized as L0==L1 using BIC were also categorized as L0==L1 using cvR$^2$ while ~200 were categorized as L0&L1 using cvR$^2$ (left most panel.) This pattern is consistent with each matrix having an upper-left triangular structure demonstrating that cvR$^2$ has a greater tendency to pick models with distinct parameters across formats (e.g. L0&L1). **(C)** Instructive example neurons illustrating how BIC and cvR$^2$ capture different notions of similarity across formats. We show the tuning of 10 example units as a function of time (mean ± sem) across a pair of formats (different rows). For each example unit, the BIC measure labeled the pairwise comparison as having a matched pattern of selectivity while the cv$R^2$ measure indicated significant but mismatched patterns of selectivity. The BIC measure captures a more qualitative notion of similarity between some (but not necessarily all) of the tested actions.

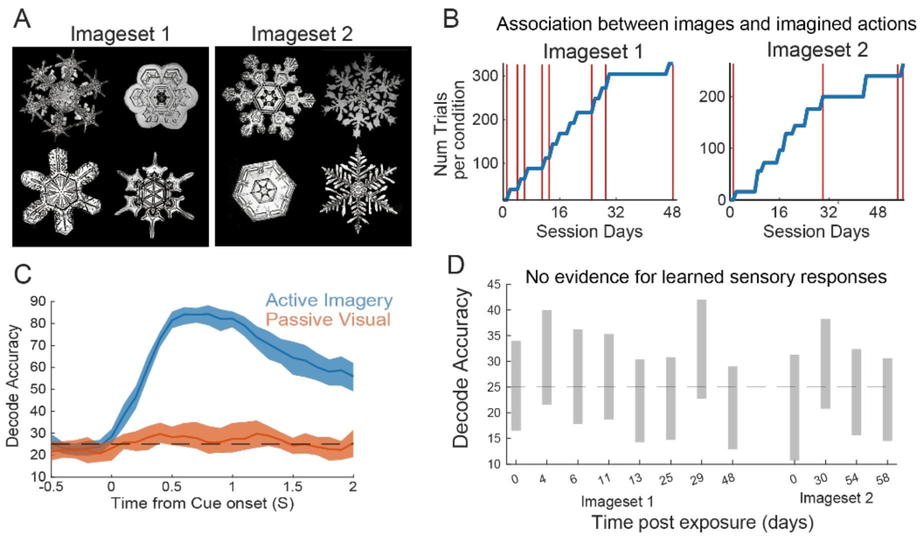


**Figure S7: Abstract symbols systematically associated with motor imagery does not evoke differential neural tuning. (A)** Abstract geometric stimuli systemtically associated with visual imagery of motor actions. **(B)** Time course of assoiative training during which symbols were used to cue imagery of motor actions (blue) and time of passive viewing of the same images (red lines). **(C)** Cross-validated classifcation of neural responses (mean ± 95% CI) during block where the subject was asked to either passively view the symbolic images (orange), or activiely perform visual imagery of motor actions upon viewing the images(blue). Horizontal dashed line indicates chance level. **(D)** Cross-validated classifcation (mean ± 95% CI) on a per-session basis for passive viewing (dashed line indicates chance level).
